# The role of H3K36 methylation and associated methyltransferases in chromosome-specific gene regulation

**DOI:** 10.1101/2021.03.04.433843

**Authors:** Henrik Lindehell, Alexander Glotov, Eshagh Dorafshan, Yuri B. Schwartz, Jan Larsson

**Author notes:** **Corresponding author**: Jan Larsson, Department of Molecular Biology, Umeå University, SE- 90187 Umeå, Sweden. **Data deposition footnote**: The RNA-seq data reported in this paper have been deposited in the Gene Expression Omnibus database (GSE166934).

## Abstract

In *Drosophila*, two chromosomes require special mechanisms to balance their transcriptional output to the rest of the genome. These are the male-specific lethal complex targeting the male X-chromosome, and Painting of fourth targeting chromosome 4. The two systems are evolutionarily linked to dosage compensation of the X-chromosome and the chromosomes involved display specific chromatin structures. Here we explore the role of histone H3 tri-methylated at lysine 36 (H3K36me3) and the associated methyltransferases in these two chromosome-specific systems.

We show that the loss of *Set2* impairs the MSL complex mediated dosage compensation; however, the effect is not recapitulated by H3K36 replacement and indicates an alternative target of Set2. Unexpectedly, balanced transcriptional output from the 4^th^ chromosome requires intact H3K36 and depends on the additive functions of NSD and the Trithorax group protein Ash1.

We conclude that H3K36 methylation and the associated methyltransferases are important factors to balance transcriptional output of the male X-chromosome and the 4^th^ chromosome. Furthermore, our study highlights the pleiotropic effects of these enzymes.

## Introduction

The evolution of sex-chromosomes, for example the X and Y chromosome pairs, often leads to sex differences in gene dose. In *Drosophila*, the Y chromosome has degenerated when it comes to gene functions, but is retained through its specific roles in male fertility (Bachtrog 2013). This leads to a difference in X-chromosome gene doses in males and females. Although some genes located on the X-chromosome are expressed in a sex-specific manner, most genes require equal expression levels in males and females (Stenberg and Larsson 2011; Mank 2013). The sex differences in gene doses of X-chromosomal genes have resulted in a need to equalize gene expression between the single X in males and the two X-chromosomes in females, and to balance this level of expression with that of the two sets of autosomal chromosomes (Oliver 2007; Stenberg and Larsson 2011; Mank 2013; Kuroda et al. 2016). In *Drosophila*, the gene dosage problem has been solved such that X-chromosomal gene expression is increased by a factor of approximately two in males (Kuroda et al. 2016). This increased expression is partly mediated by chromosome-specific targeting and stimulation of the male X-chromosome by the male-specific lethal (MSL) complex. The MSL complex consists of at least five protein components, originally identified through their male-specific lethal phenotype when lost; MSL1, MSL2, MSL3, MLE and MOF, and two long non-coding RNAs (lncRNAs) named *RNA on the X* (*roX1* and *roX2*) (Kuroda et al. 2016; Samata and Akhtar 2018). The targeting of the MSL complex to the male X-chromosome is in principle described in two steps. The MSL complex initially binds specifically to roughly 250 specific sites, denoted: chromatin entry sites (CES); MSL recognition element (MRE); high-affinity sites (HAS); or pioneering sites on the X (Pion-X), largely overlapping but initially isolated and classified using different criteria (Kelley et al. 1999; Alekseyenko et al. 2008; Straub et al. 2008; Villa et al. 2016). Next, the MSL complex spreads from these high-affinity sites to neighbouring transcriptionally active genes; this spreading requires the presence of at least one of the two *roX* RNAs (Figueiredo et al. 2014). The resulting X-chromosome specific targeting is accompanied by an enriched histone 4 lysine 16 acetylation (H4K16ac), mediated by the acetyltransferase MOF (Akhtar and Becker 2000; Smith et al. 2000).

Importantly, at least one more chromosome has been the subject of as similar an evolutionary process as the current X-chromosome. The 4^th^ chromosome in *Drosophila melanogaster* was ancestrally an X-chromosome that has reverted to an autosome (Vicoso and Bachtrog 2013; Vicoso and Bachtrog 2015). This sex-chromosome reversal is likely to explain that the 4^th^ chromosome in *D. melanogaster* is the subject of chromosome-specific targeting and regulatory mechanisms mediated by the protein Painting of fourth (POF) (Larsson et al. 2001; Larsson et al. 2004; Johansson et al. 2007a; Kim et al. 2018a). We have previously hypothesised that POF and its stimulatory function became trapped on the 4^th^ chromosome when the latter reverted to being an autosome (Kim et al. 2018a). The stimulatory effect on transcript output mediated by POF is comparable in level to the compensation mediated by the MSL complex on the X-chromosome. The hypothesised origin of POF as a dosage compensation system is also supported by the fact that the POF mediated compensation of the 4^th^ chromosome is essential for the survival of flies with a monosomy of chromosome 4 (Johansson et al. 2007a). The 4^th^ chromosome, targeted by POF, has several unique characteristics. It is the smallest chromosome in the *Drosophila* genome, it is replicated late, and in principle, the entire 4^th^ chromosome can be considered heterochromatic (Riddle and Elgin 2006; Filion et al. 2010). This means that the 4^th^ chromosome is enriched in HP1a, the histone methyltransferase Setdb1, as well as methylated H3K9 (Riddle et al. 2011; Figueiredo et al. 2012).

Accurate targeting and the precise stimulatory effect of these chromosome-specific systems require the contributions of several factors to be well coordinated. In addition to specific DNA sequence motifs mentioned above, these may include distinct DNA topology and spatial positioning of the dosage-compensated chromosomes (Samata and Akhtar 2018; Valsecchi et al. 2020) as well as the interaction between the chromo-domain of MSL3 and histone H3 tri-methylated at lysine 36 (H3K36me3). It was reported that in *Set2*^*1*^ mutants, with reduced levels of H3K36me3, the localization of MSL2 and MSL3 toward gene ends was impaired (Larschan et al. 2007) and that the interaction between MSL3 and H3K36me3 helps the MSL complex to bind active genes (Larschan et al. 2007; Sural et al. 2008). However, structural studies suggest that the chromo-domain of MSL3 has higher affinity to the more abundant histone H4 mono- and di-methylated at lysine 20 (H4K20) (Kim et al. 2010; Moore et al. 2010); the rival hypothesis contends that the MSL3 chromo-domain interacts with methylated H4K20 to present H4 tails for acetylation at lysine 16 by MOF (Moore et al. 2010). Therefore, the extent and mechanisms by which H3K36 methylation and/or associated methyltransferases contribute to balanced transcriptional output from the male X and the 4^th^ chromosomes remains an open question.

Three evolutionarily conserved *Drosophila* proteins: Absent, small or homeotic discs 1 (Ash1), SET domain containing 2 (Set2) and Nuclear receptor binding SET domain containing protein (NSD), are believed to methylate H3K36. *In vitro*, Ash1 can add one or two methyl groups to H3K36 (Tanaka et al. 2007) and this activity is further enhanced by Mrg15 and Caf1 proteins with which Ash1 forms a complex (Huang et al. 2017; Schmahling et al. 2018; Hou et al. 2019; Lee et al. 2019). Flies lacking *ash1* function show an approximated two-fold reduction of bulk H3K36me1 but no detectable loss of overall H3K36me2 or H3K36me3 (Schmahling et al. 2018; Dorafshan et al. 2019). These *ash1* mutants display multiple homeotic transformations and die at the larval stage due to erroneous repression of developmental genes by Polycomb group mechanisms (Tripoulas et al. 1994; Klymenko and Muller 2004; Dorafshan et al. 2019). The Set2 protein can methylate H3K36 *in vitro* using mono- and di-methylated H3K36 as a substrate (Bell et al. 2007). Whether it can also add methyl groups to unmethylated H3K36 is not entirely clear although it has been demonstrated that the human orthologue (SETD2) is able to do so in reconstituted reactions (Yuan et al. 2009). Consistently, the *Drosophila Set2* mutants have 10-fold lower levels of H3K36me3 overall and at specific genes (Larschan et al. 2007; Dorafshan et al. 2019) but display no major changes in the bulk H3K36me2 and H3K36me1 (Dorafshan et al. 2019). Taken together, these observations suggest that Set2 produces most of the *Drosophila* H3K36me3 and may contribute to cellular pools of H3K36me1 and H3K36me2. The loss of *Set2* function is lethal (Larschan et al. 2007; Dorafshan et al. 2019), but the corresponding mutant shows no signs of excessive Polycomb repression (Dorafshan et al. 2019) suggesting that Set2 and Ash1 affect *Drosophila* development in distinct ways. Less is known about the biochemical properties of the fly NSD. It has three closely related orthologues in mammals (NSD1, NSD2, NSD3) whose methyltransferase activity *in vitro* has been extensively studied. These studies concur that NSD proteins can mono- and di-methylate H3K36 (Rayasam et al. 2003; Li et al. 2009; Kudithipudi et al. 2014). However, methylation of other substrates is also suggested. A systematic screen indicates that these include K168 of histones H1.5 and H1.2, K169 of histone H1.3, K44 of histone H4 as well as K1033 of the chromatin re-modeller ATRX and K189 of the U3 small nucleolar RNA-associated protein 11 (Kudithipudi et al. 2014). NSD proteins are essential for proper mammalian development and haploinsufficiencies in the *NSD1* and *NSD2* genes were linked to Sotos and Wolf-Hirschhorn genetic syndromes, respectively (Kurotaki et al. 2002). Whether methylation of H3K36 or any of the substrates above are relevant for the developmental functions of the NSD proteins remains an open question. Unlike its mammalian counterparts, *Drosophila NSD* loss of function mutants are viable, fertile, and show no obvious morphological defects (Dorafshan et al. 2019). Although early experiments in cultured cells suggested that NSD knock-down reduces bulk levels of H3K36me2 and H3K36me3 (Bell et al. 2007), the *NSD* loss of function mutants display no obvious changes in the overall levels of H3K36me1, H3K36me2 and H3K36me3 (Dorafshan et al. 2019).

Whether H3K36 is the only (or even the major) physiological substrate is a question that appears equally relevant for Ash1 and Set2. One might expect that animals in which H3K36 is replaced with an amino acid that cannot be methylated, will display defects similar to those seen in mutants that lack the corresponding methyltransferase. Mounting experimental evidence suggests the contrary. Thus complete substitution of the zygotic H3 with a variant in which K36 is replaced with arginine (R) does not cause excessive repression of homeotic genes seen in *ash1* mutants (Dorafshan et al. 2019). Likewise, flies with replication-coupled histone H3.2 replaced with the H3K36R variant, display no cryptic transcription initiation or altered splice site choice as reported for yeast *Set2* or human *SETD2* mutants (Meers et al. 2017).

To investigate the role of H3K36 and the associated histone methyltransferases in chromosome-specific gene regulation we evaluated relative levels of methylated H3K36 on individual chromosome arms and compared those to transcriptional imbalance caused by the loss of Set2, NSD and Ash1 or the replacement of the histone H3.2 with a variant in which lysine 36 is substituted to arginine.

Confirming previous reports, we found that loss of *Set2* impairs MSL complex mediated dosage compensation. However, the effect is not recapitulated in H3K36R mutants and suggests an alternative target for Set2. This implies that the model in which Set2-mediated H3K36me3 helps the MSL complex to bind active genes may need to be revised. Unexpectedly, our results indicate that the balanced transcriptional output from the 4^th^ chromosome depends on the additive contribution from Ash1 and NSD and requires intact H3K36. We conclude that H3K36 methylation and the associated methyltransferases are important factors for balanced transcriptional output of the male X-chromosome and the 4^th^ chromosome. Our study also emphasises the importance of pleiotropic effects of these enzymes.

## Results

### H3K36me3 is reduced on the male X-chromosome and enriched on the 4^th^ chromosome

*Drosophila* chromosome-specific regulatory systems are accompanied by chromosome-specific enrichment of specific histone modifications. The male X-chromosome is enriched in H4K16ac (Turner et al. 1992) and the 4^th^ chromosome is enriched in H3K9me2/me3 (Czermin et al. 2002). Considering the proposed role of methylated H3K36 in dosage compensation, we decided to analyse methylation of H3K36 and the associated methyltransferases further. Immunostaining of polytene chromosomes from male third instar larvae with antibodies against mono- di- and tri-methylated H3K36 shows that these modifications are differentially distributed between the chromosome arms. Mono-methylated H3K36 (H3K36me1) was evenly distributed in a banded pattern throughout the entire genome (Supplementary Fig.S1A). Di-methylated H3K36 (H3K36me2) was also ubiquitous genome-wide, but the banding pattern of H3K36me2 shows only minor overlaps with H3K36me1 (Supplementary Figs. S1A, B). A noticeable difference is the increased enrichment of H3K36me2 in the pericentromeric heterochromatin as well as on the heterochromatic fourth chromosome compared to the major chromosome arms (Supplementary Fig. S1A). Increased enrichment on the fourth chromosome was also consistently observed for H3K36me3 (Fig. 1A) but unlike H3K36me2, less relative enrichment in pericentromeric heterochromatin was observed for H3K36me3 (Supplementary Fig. S1C). Unexpectedly, H3K36me3 appeared less abundant on the male X-chromosome (Fig. 1A, Supplementary Fig. S1C) compared to the autosomes. Importantly, weaker anti-H3K36me3 immunostaining of the X-chromosome was not observed in females (Fig. 1B). In males, the number of chromatids of the polytene X-chromosome is half that of the autosomes. This difference may account for some of the reduced staining observed. Nevertheless, the decrease in H3K36me3 staining is still more pronounced than that seen for H3K36me1 (Supplementary Fig. S1A) or H3K36me2 (Supplementary Fig. S1C). This argues that the male polytene X-chromosome has a lower level of H3K36me3 than the autosomes. The apparent lower level of H3K36me3 on the male X-chromosome, together with its enrichment on the fourth chromosome, is at odds with the model in which Set2-mediated H3K36me3 helps the MSL complex to bind active genes (Larschan et al. 2007; Sural et al. 2008). We, therefore, investigated the question further.

**Figure 1.**
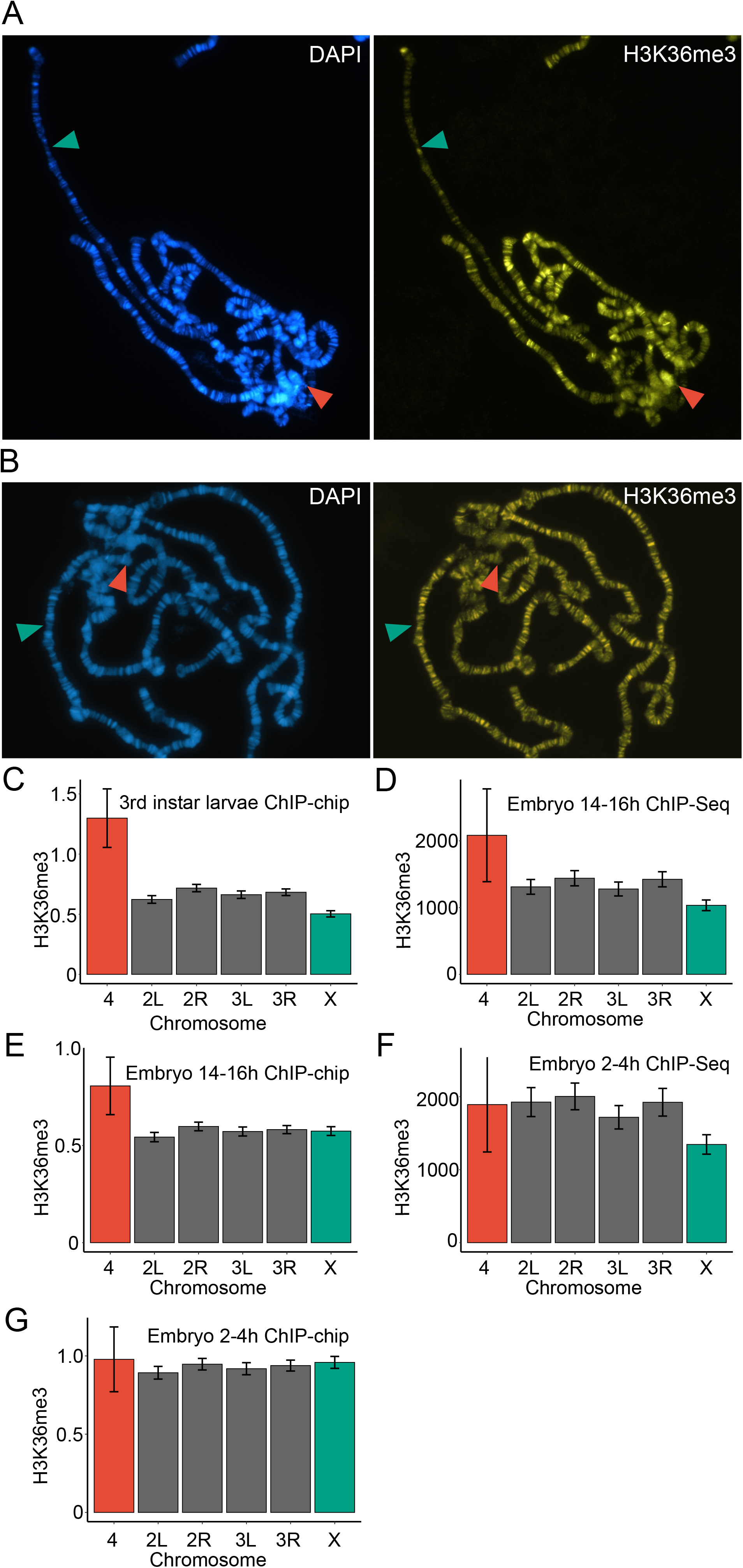
H3K36 tri-methylation is enriched on the 4^th^ chromosome and reduced on the male X-chromosome. (*A*) Immunostaining of a male third instar larvae polytene chromosome shows an accumulation of H3K36me3 on chromosome 4 (red arrow) and a reduction on the X-chromosome (green arrow) as compared to autosomal signals. H3K36me3 in yellow and DAPI staining of DNA in blue. (*B*) Immunostaining of female third instar larvae polytene chromosome with no observable differences between any chromosome arms. H3K36me3 in red and DAPI staining in blue. The 4^th^ chromosome and the X-chromosome are indicated with red and green arrows, respectively. (*C*) Average exon H3K36me3 ChIP-chip enrichment scores per chromosome in mixed-sex third instar larvae. This ChIP data analysis confirms the observations from the immunostainings that H3K36me3 is significantly enriched on the 4^th^ chromosome and likewise reduced on the male X-chromosome. (*D*) Average exon H3K36me3 ChIP-Seq enrichment scores per chromosome in mixed-sex late embryos (14 – 16 h). Chromosome 4 is significantly more enriched in H3K36me3 while the X-chromosome is reduced in H3K36me3 as compared to the major autosome arms. (*E*) Average exon H3K36me3 ChIP-chip enrichment scores per chromosome in mixed-sex late embryos (14 – 16 h). Chromosome 4 is significantly more enriched in H3K36me3 compared to the autosomes. (*F, G*) Average exon H3K36me3 ChIP-Seq enrichment scores per chromosome in mixed-sex early embryos (2 – 4 h). The ChIP-Seq data show lower H3K36me3 levels on the X-chromosome (*F*). No significant differences were observed in ChIP-chip from early embryos (*G*). Enrichment scores for the ChIP-chip experiments (*C, E, G*) is the ChIP over input log_2_ ratio of H3K36me3 in the top 50% of the exon regions per gene. For ChIP-seq (*D, F*) the H3K36me3 enrichment scores are reads adjusted for difference in position between ChIP and input. In ChIP-seq only peaks in exons were used. Error bars (*C - G*) indicate the 95% confidence intervals. Chromosome 4 is shown in red, autosomes in grey and the X-chromosome is shown in green.

### Chromosome-specific differences in tri-methylated H3K36 are established during early development

To validate and quantify our observations in polytene tissue, we analysed H3K36me3 ChIP-chip data from third instar *D. melanogaster* larvae (mixed sexes) available from modENCODE (mod et al. 2010) as well as the more recent H3K36me3 ChIP-seq data (Prayitno et al. 2019). Since H3K36me3 is predominantly enriched within the coding regions of genes (Schwartz et al. 2009), we calculated relative enrichment values of H3K36me3 within exons for each gene. The average ChIP signals divided by chromosomes confirmed our observation from the polytene chromosome stainings. We found that H3K36me3 ChIP signals within gene bodies in 3^rd^ instar larvae were higher on the fourth chromosome and lower on the male X-chromosome as compared to the autosomes (Fig. 1C). Analysis of H3K36me3 in late embryonic stage (mixed sexes) ChIP-Seq data (Prayitno et al. 2019) showed the same distribution pattern (Fig. 1D).

Next, we analysed the H3K36me3 abundance over developmental time by looking at additional ChIP datasets from different developmental stages (mod et al. 2010; Prayitno et al. 2019). ChIP-chip data in early embryos (2 – 4 h) indicate that the amount of H3K36me3 is uniformly the same for all chromosomes (Fig. 1G). In contrast, the ChIP-seq data show reduced H3K36me3 ChIP signal on the X-chromosome, even at this early stage (Fig. 1F). In a late embryonic stage (14 - 16 h), the fourth chromosome shows significantly stronger immunoprecipitation with H3K36me3 antibodies in both ChIP assays (Figs. 1D, E). However, only the ChIP-seq data show a reduced ChIP signal on the X-chromosome (Fig. 1D). By the 3^rd^ instar larval stage, as shown in Fig. 1C, the ChIP-signal on the X-chromosome is lower than on the autosomes and the signal on the fourth is even more pronounced.

### The level of H3K36me3 within genes correlates with the degree of transcriptional activity and gene length

We and others have shown that the two chromosome-specific systems studied here affect genes differently depending on gene type, expression levels, gene length and replication timing (Schubeler et al. 2002; Gilfillan et al. 2006; Stenberg et al. 2009; Regnard et al. 2011; Lundberg et al. 2013b; Philip and Stenberg 2013; Lubelsky et al. 2014; Lee et al. 2016; Kim et al. 2018b; Prayitno et al. 2019; Belyi et al. 2020; Ekhteraei-Tousi et al. 2020). We therefore determined how these criteria relate to the levels of H3K36me3 within genes. We found that the H3K36me3 ChIP signal at the gene level positively correlates with wildtype transcript abundance (Fig. 2A) and longer genes accumulate more H3K36me3 compared to short genes (Fig. 2B). We also observed higher enrichment of H3K36me3 in early replicating genes, compared to late replicating genes, on all chromosomes (Fig. 2C). Genome-wide housekeeping genes were also significantly more enriched by H3K36me3 than non-housekeeping genes (Fig. 2D).

**Figure 2.**
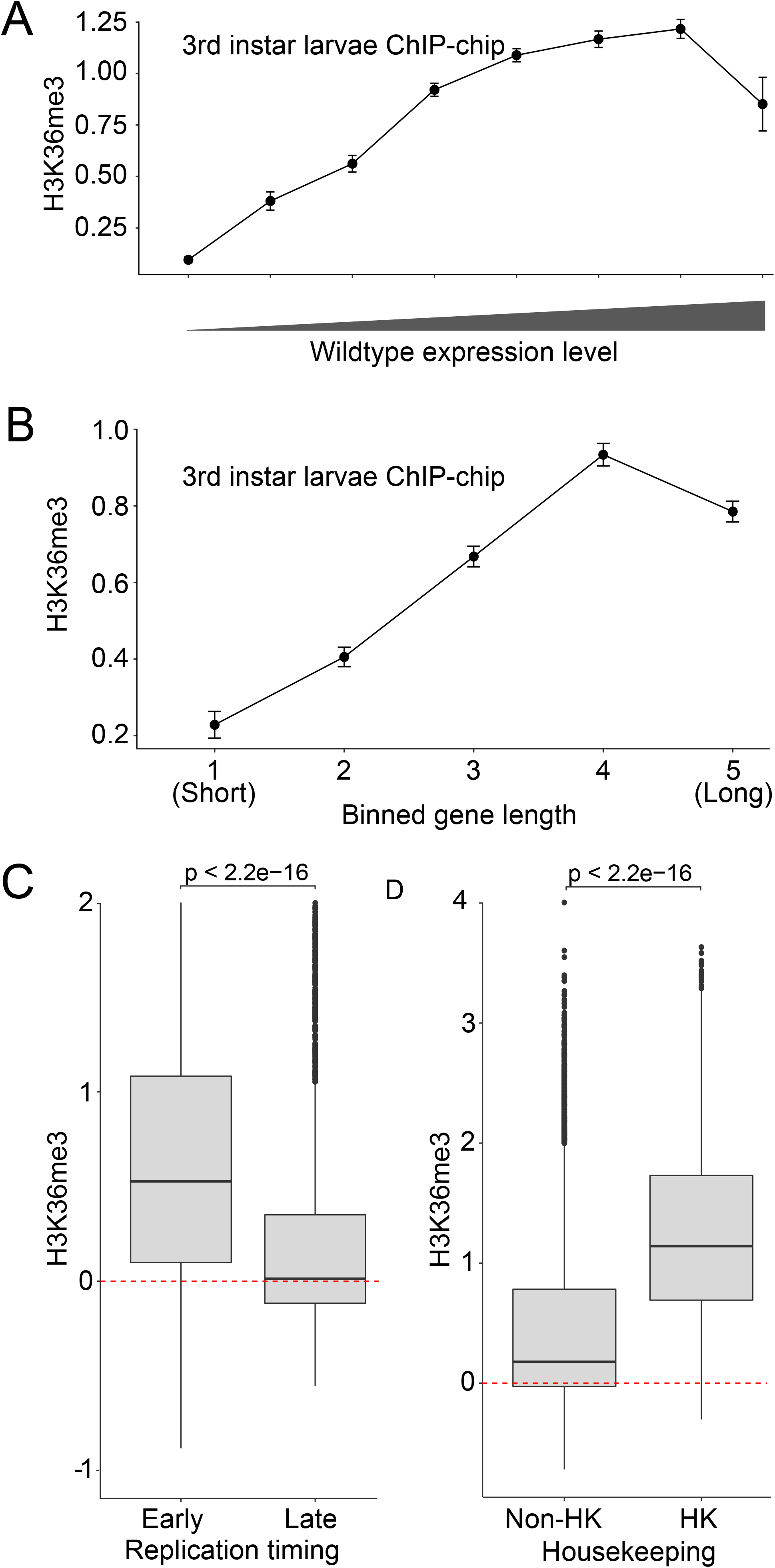
Gene levels of H3K36me3 ChIP signal positively correlate with gene lengths and expression levels. Third instar larvae ChIP-chip gene exon enrichment level (ChIP over input, log2) plotted against: (*A*) Transcript abundance in wildtype male third instar larval brains as determined by the RNA-Seq experiment. Highly transcribed genes show a higher accumulation of H3K36me3 than do genes with lower transcript abundance. (*B*) Gene length, i.e., the total gene length (5′-3′) divided into five equal sized bins. Longer genes accumulate more H3K36me3. (*C*) Early replicating genes accumulate more H3K36me3 than late replicating genes. (*D*) Housekeeping genes accumulate more H3K36me3 than do non- housekeeping genes. H3K36me3 ChIP-chip enrichment scores represent the ChIP over input log2 ratio of H3K36me3 in the top 50% of the exonic regions per gene. In (*A, B*) the error bars indicate the 95% confidence intervals. For (*C, D*) the statistical significances were determined by Unpaired Two-Sample Wilcoxon Tests.

#### Set2 is required for balanced transcription of X-linked genes bound by the MSL complex

The level of H3K36me3 within genes correlates with their transcriptional activity, and Set2 has been reported to help in maintaining the transcriptional balance of the male X-chromosome. Yet the level of H3K36me3 on this chromosome appears to be lower than it is on autosomes. Puzzled by this paradox, we decided to analyse transcriptional outputs from each chromosome in mutants of the three known H3K36 specific methyltransferases: Set2, NSD and Ash1. In flies, Set2 is responsible for most of the H3K36me3 while the potential division of labour and redundancy in H3K36 methylation mediated by NSD and Ash1 is not fully understood. As reported previously (Alekseyenko et al. 2014), we found that NSD is enriched in pericentromeric heterochromatin and on the 4^th^ chromosome on polytene chromosomes (Supplementary Fig. S2), with a staining pattern similar to HP1a and to the H3K36me2 enrichment (Supplementary Fig. S1B) (James et al. 1989; Figueiredo et al. 2012; Alekseyenko et al. 2014).

It has been reported that in *Set2*^*1*^ mutants the MSL complex correctly targets the male X-chromosome but with reduced binding levels to its target genes (Larschan et al. 2007). The *Set2*^*1*^ allele corresponds to a deletion of the N-terminal half of its open reading frame, including the catalytic SET domain (Larschan et al. 2007) and leads to approximately a 10- fold reduction in global H3K36me3 (Larschan et al. 2007; Dorafshan et al. 2019). Homozygous *Set2*^*1*^, but not *ash1*^*22*^*/ash1*^*9011*^ or *NSD*^*ds46*^ (see below), displays a significant relative decrease in transcriptional output from the male X-chromosome (Fig. 3A) and a nearly complete loss of H3K36me3 staining on polytene chromosomes (Supplementary Fig. S3). By dividing genes according to their wildtype transcript levels we found that highly transcribed genes are those most affected by loss of Set2 (Fig. 3B). We have previously shown that highly transcribed genes require high levels of MSL complex binding (Kim et al. 2018b). We therefore asked if the reduced expression observed in *Set2*^*1*^ correlates with binding levels of the MSL complex. To address this question, all X-chromosome genes were divided into five bins based on their immunoprecipitation with antibodies against the MSL1 subunit of the MSL complex (Kind et al. 2008; Philip and Stenberg 2013). Thus, bin 1 included unbound and weakly bound genes, while bin 5 included genes highly enriched in MSL proteins. As illustrated in Fig. 3C, genes binding MSL complex tend to show reduced transcript abundance upon Set2 loss. To further substantiate this result we grouped the genes by distance to the nearest High Affinity Site (HAS) since it has been shown that genes more distal to HAS are less sensitive to loss of a functional MSL complex (Straub et al. 2008; Kim et al. 2018b). As expected, we observed that genes close to a HAS are, on average, more affected by the loss of Set2 compared to genes further away. Genes more distal than about 30 kb from a HAS are essentially unaffected by the loss of Set2 (Fig. 3D). This is in line with what we have observed after ablation of the *roX1 roX2* non-coding RNAs (Kim et al. 2018b). We conclude that the reduced transcript levels on the male X-chromosome observed in *Set2*^*1*^ corresponds well to a dysfunction in MSL complex mediated dosage compensation.

**Figure 3.**
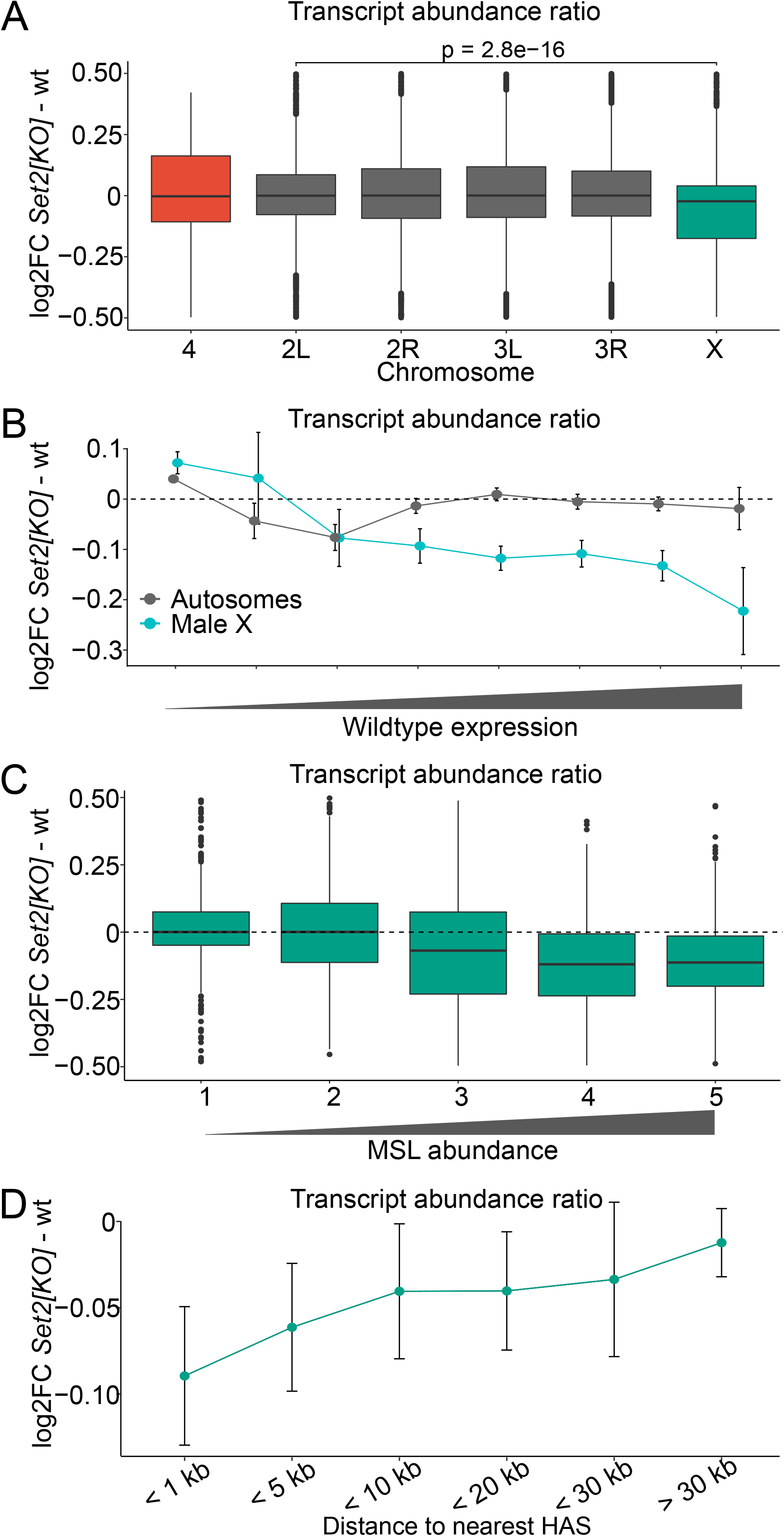
*Set2* is required for balanced transcription output from the single male X- chromosome. (*A*) Boxplot showing chromosome-specific transcript abundance ratios (log_2_ fold change) between *Set2*^*1*^ and wildtype for chromosome 4 (red), X-chromosome (green) and the autosome arms (grey). The transcript abundance is significantly reduced for the X- chromosomes as compared to the autosomes in *Set2*^*1*^. Significance levels were determined by Unpaired Two-Sample Wilcoxon Tests between the X-chromosome and chromosome arm2L (the least unlike autosome arm). (*B*) Transcript abundance in *Set2*^*1*^ versus wildtype plotted as fold change ratio (log_2_). Genes were binned according to Flybase RNA-Seq expression level intervals. Genes with higher wildtype expression output are significantly more affected on the X-chromosome than on the autosomes. (*C*) Boxplot showing average transcript abundances of the X-chromosomal genes in *Set2*^*1*^ versus wildtype. The genes are grouped in equally sized bins based on their MSL1 binding strength; 1 (lowest) to 5 (highest). Genes with higher MSL1 levels show reduced transcript abundance compared to genes with no or low amounts of MSL1 bound. (*D*) Average log_2_ fold change for X-chromosomal genes binned by distance to high-affinity sites (HAS). In (*B*) and (*D*) the error bars indicate the 95% confidence intervals.

### H3K36R mutants show no imbalance of transcriptional output from the male X-chromosome

Loss of *Set2* function has been shown to cause reduced MSL complex binding to target genes (Larschan et al. 2007) and our results indicate that the *Set2*^*1*^ mutation causes a relative decrease in transcriptional output from the male X-chromosome in a pattern that mimics impaired function of the MSL complex. It is tempting to speculate that the loss of Set2 leads to lower H3K36me3 which, in turn, impairs the binding of MSL3 (Larschan et al. 2007; Sural et al. 2008). However, the interaction between H3K36me3 and the chromo-domain of MSL3, proposed for this mechanism, is at odds with structural data (Kim et al. 2010; Moore et al. 2010) and our previous proximity ligation analysis did not detect a direct interaction between MSL3 and H3K36me3 (Lindehell et al. 2015). If the reduction of the transcriptional output from the male X-chromosome observed in the *Set2*^*1*^ mutants is caused by a concomitant reduction in H3K36me3, we would expect to see a similar transcriptional imbalance in the H3K36R histone replacement mutant, *ΔHisC; 12x*^*H3K36R*^. These mutant flies carry the deletion of the histone gene cluster *ΔHisC* combined with a transgenic construct carrying twelve copies of the 5 kb histone repeat unit in which H3.2K36 is mutated to arginine (*12x*^*H3K36R*^) (Gunesdogan et al. 2010; McKay et al. 2015).

Polytene chromosomes of the *ΔHisC; 12x*^*H3K36R*^ flies display a dramatic decrease of immunostaining with antibodies against H3K36me3 (Fig. 4A). This is in accordance to what has previously been shown (McKay et al. 2015) and is similar to that seen in *Set2*^*1*^ mutants (Supplementary Fig. S3). Western blot analysis showed that H3K36me3 methylation is, as expected, strongly reduced in *ΔHisC; 12x*^*H3K36R*^ flies (Fig. 4B). Notably, some H3K36me3 signal is still detected. This may correspond to the tri-methylated H3.3 variant histone still present in *ΔHisC; 12x*^*H3K36R*^ flies; but we cannot exclude the possibility that some of it may be caused by antibody cross-reactivity. Despite the five-fold reduction in the overall H3K36me3, we observed no X-chromosome specific transcriptional imbalance in *ΔHisC; 12x*^*H3K36R*^ animals (Fig. 4C), which suggests that imbalance in transcription of genes on male X-chromosome after Set2 knock-out is not linked to a concomitant loss of H3K36 methylation. The interpretation of histone replacement experiments may, however, be less straightforward if it is the un-methylated H3K36 that is required for the process, which Set2 may aim to prevent. Thus, recent structural studies suggest that unmodified H3K36 is required for unimpeded methylation of lysine 27 of histone H3 (H3K27) by the Polycomb Repressive Complex 2 (PRC2) (Jani et al. 2019; Finogenova et al. 2020). Consistently, flies in which the zygotic histone H3.2 is replaced with a variant that contains arginine instead of K36, show mild defects in Polycomb repression of homeotic selector genes (Finogenova et al. 2020). We, therefore, considered a possibility that reduced activity of PRC2 in the *ΔHisC; 12x*^*H3K36R*^ mutant may lead to an increased transcription of the X-chromosome genes, which in turn, may mask the effect of the Set2 knock-out. To address this issue, we took advantage of the RNA-seq data from (Lee et al. 2015) where the PRC2 function was disrupted by a temperature shift of a cell line homozygous for the temperature sensitive *E(z)*^*61*^ allele (Jones and Gelbart 1990; Lee et al. 2015). We calculated transcript abundance ratios for all genes divided by chromosomes and found the transcription of genes on the X-chromosome relative to autosomes to be reduced, not increased (Fig. 4D). Whether this relative reduction is a consequence of global de-repression of low and non-expressed genes as noted previously (Lee et al. 2015) remains an open question which we have not attempted to investigate further at this stage. Nevertheless, we find no evidence that the lack of an X-chromosome specific effect upon H3K36R replacement is caused by potential crosstalk with PRC2. Taken together, our results argue that the reduced transcription of dosage compensated genes after Set2 knock-out is not linked to a concomitant loss of H3K36 methylation.

**Figure 4.**
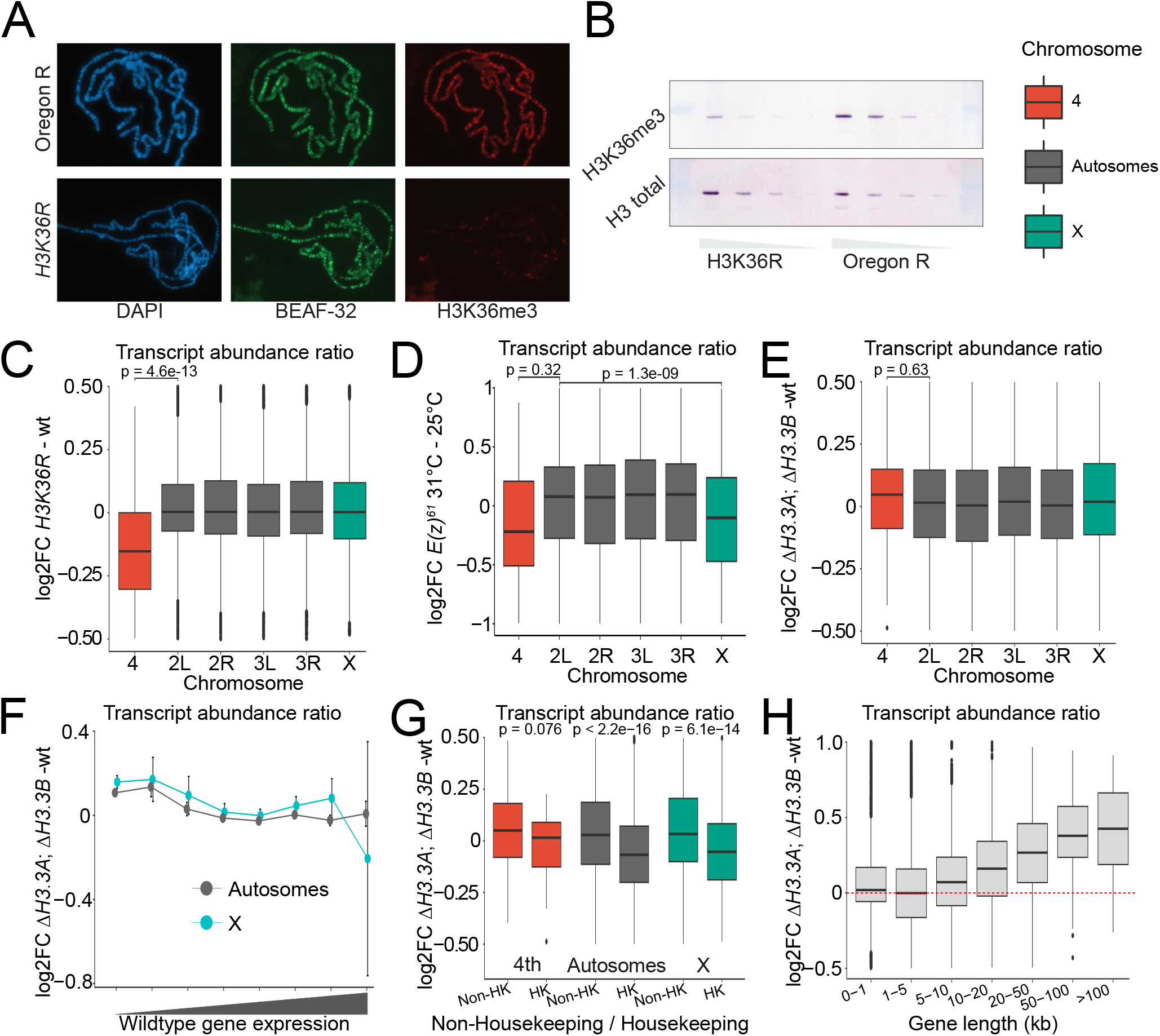
The replication independent histone variant H3.3 is required for proper transcription of short, moderately to high expressed genes. (*A*) Staining of H3K36me3 in salivary glands for wildtype (top row) and *ΔHisC; 12x*^*H3K36R*^ histone replacement (bottom row). Staining of BEAF-32 serves as an internal control. (*B*) Western blot of brains and imaginal discs of third instar larvae showing significant decrease of H3K36me3 (top row) in *ΔHisC; 12x*^*H3K36R*^ versus wildtype. Total Histone 3 levels serve as loading control (2-fold dilutions steps). (*C*) Boxplot showing log_2_ fold change of transcript abundance per chromosome for *ΔHisC; 12x*^*H3K36R*^ versus *ΔHisC; 12x*^*H3K36K*^ (wt). Note that chromosome 4 shows a significant reduction in transcript abundance compared to the other chromosome arms. (*D*) Box plot showing log_2_ fold change of transcript abundance per chromosome for the *E(z)*^*61*^ cell line at 31°C (strongly reduced E(z) function) versus 25°C). The transcript abundance is significantly reduced for the X-chromosomes as compared to the major autosome arms upon reduced E(z) function. (*E*) Boxplot showing log_2_ fold change of transcript abundance per chromosome for *ΔH3*.*3B; ΔH3*.*3A* versus wildtype flies. No significant differences are observed between chromosomes. (*F*) Transcript abundance ratio (log_2_ fold change) in *ΔH3*.*3B; ΔH3*.*3A* versus wildtype in relation to gene expression levels in wildtype. Genes were binned based on wildtype expression output level. Genes on the X-chromosome, as well as genes on autosomes with moderate to high expression levels, were more sensitive to the loss of H3.3. (*G*) Boxplot showing log_2_ fold change of transcript abundance in *ΔH3*.*3B; ΔH3*.*3A* compared to wildtype with genes divided into non-housekeeping (Non-HK) and housekeeping (HK). Genes on chromosome 4 and the X-chromosome are indicated (4^th^) and (X), respectively. (*H*) Boxplot showing log_2_ fold change of transcript abundance in *ΔH3*.*3B; ΔH3*.*3A* compared to wildtype, genome-wide. Genes were binned by total gene lengths. The increase in transcript abundance in *ΔH3*.*3B; ΔH3*.*3A* compared to wildtype positively correlates with gene length. For (*C - F*) significant differences were determined by Unpaired Two-Sample Wilcoxon Tests. Error bars in (*F*) indicate 95% confidence intervals.

We next considered the possibility that methylation of variant histone H3 (H3.3) is critical for balanced transcriptional output from the male X-chromosome and the regulatory mechanism mediated by Set2 methylation. It has previously been shown that the replication independent histone variant H3.3 is enriched on the male X-chromosome (Mito et al. 2005). We therefore analysed gene transcription in flies with both *H3*.*3* genes deleted, *ΔH3*.*3B; ΔH3*.*3A*. Despite the reported enrichment of H3.3 on the male X-chromosome we observed no reduction in X-chromosome transcript abundance as compared to autosomes in the *ΔH3*.*3B; ΔH3*.*3A* double mutant (Fig. 4E).

A minor chromosome-specific effect on a subset of genes could be masked when all genes are analysed together. Intriguingly, when the genes were binned into subcategories, interesting trends emerged. We noted that genes with moderate to high transcription levels were more sensitive to the loss of H3.3. However, this trend was observed on the autosomes as well as on the male X-chromosome (Fig. 4F). Likewise, the transcript levels from housekeeping genes were decreased as compared to non-housekeeping genes (Fig. 4G). We also observed a small but significant relative increase of transcript abundance from late replicating genes (Supplementary Fig. S4). The most striking effect was in relation to gene length, where we observed a clear positive correlation between the gene length and the transcriptional changes in *ΔH3*.*3B; ΔH3*.*3A* mutants (Fig. 4H). We conclude that, despite the relative enrichment of H3.3 on the male X-chromosome, the loss of H3.3 has no chromosome-specific effect measurable in our assay. However, the proper transcription of short, moderately to highly transcribed genes, is still dependent on H3.3 in a non chromosome-specific manner. Taken together, the lack of an X-chromosome specific effect in *ΔHisC; 12x*^*H3K36R*^ as well as in *ΔH3*.*3B; ΔH3*.*3A* mutants argues that Set2 contributes to increased expression of dosage compensated genes independently of H3K36 methylation.

### The 3′-UTR length correlates with transcriptional activity and amount of H3K36me3 within genes

It has been shown that H3K36me3 levels positively correlate with the length of the 3′-UTR in *Drosophila* as well as in *C. elegance* (Pu et al. 2015). In agreement with this, we observed a positive correlation of both the wildtype gene transcription (transcript abundance) and the strength of H3K36me3 ChIP signal within gene bodies to the length of the 3′-UTR (Fig. 5A, B). Importantly, when we calculated the average length of the 3′-UTR chromosome wise, we found that the 4^th^ chromosome and the X-chromosome have 2.07 and 1.46 times longer 3′- UTRs on average, compared to the autosomal average (Fig. 5C). We therefore asked whether the decrease in the transcriptional output from the X-chromosome genes in the *Set2*^*1*^ mutants differs depending on UTR length, and compared it to those in the *ΔHisC; 12x*^*H3K36R*^ and the *ΔH3*.*3B; ΔH3*.*3A* mutants (Figs. 5D-F). We found no correlation between the decrease in the transcriptional output from the X-chromosome genes in the *Set2*^*1*^ mutants and the length of their UTRs (Fig. 5D). However, genes with long UTRs showed increased transcript abundance in the *ΔH3*.*3B; ΔH3*.*3A* mutants (Fig. 5F), which is in line with the increased transcript abundance observed for long genes (Fig. 4H). We conclude that the two chromosomes with chromosome-specific gene regulatory mechanisms both have longer average 3′UTRs, a characteristic that correlates with higher transcript abundance and higher H3K36me3 ChIP signal.

**Figure 5.**
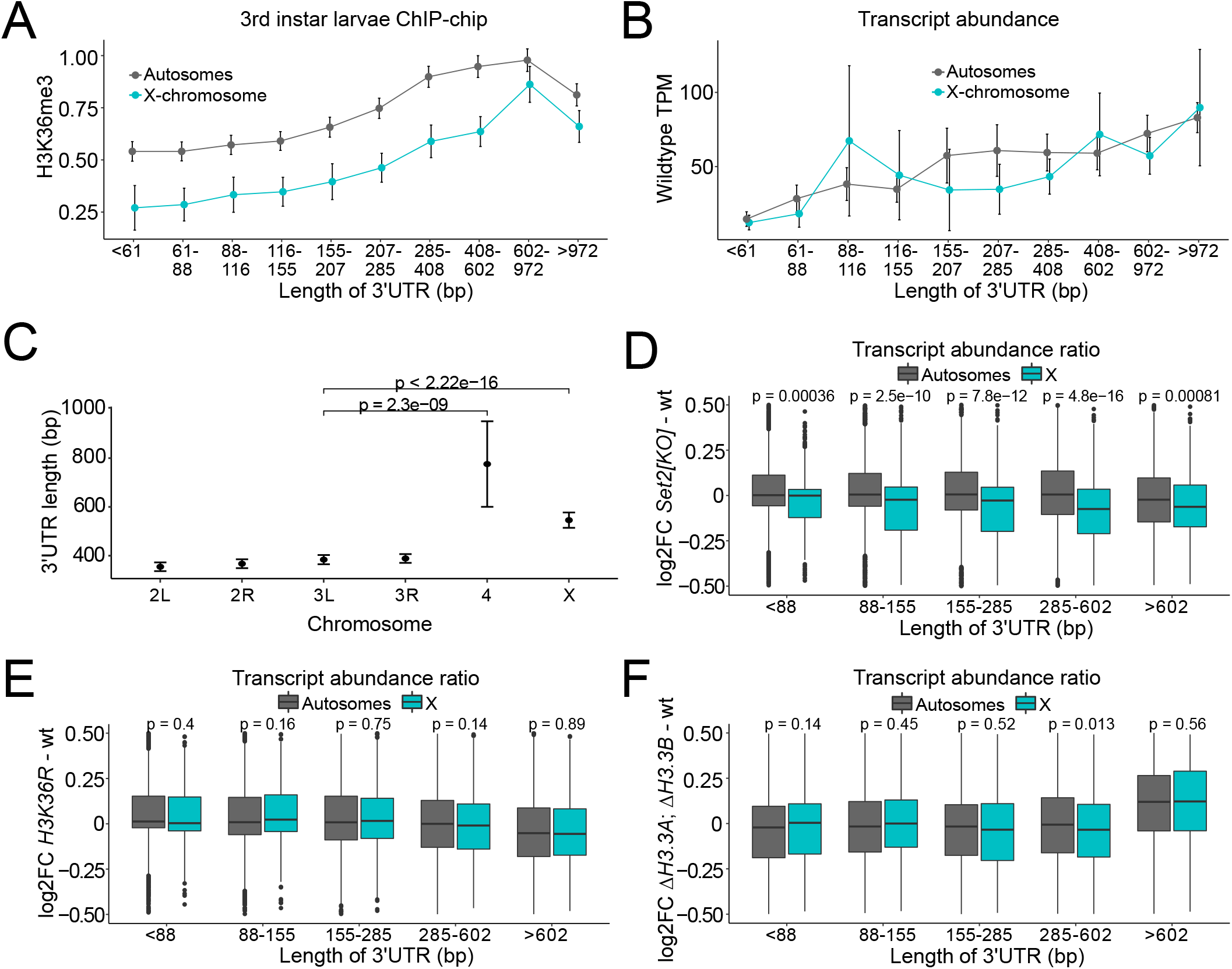
The X- and the 4^th^ chromosomes have longer 3′-UTRs, a characteristic that positively correlates with gene expression output and H3K36me3 enrichment. (*A*) Line graph showing a positive correlation between gene level enrichments of H3K36me3 and gene average 3′-UTR length for the X-chromosome (green) and the autosomes (grey). The genes are binned into equally sized groups based on their 3′-UTR length. (*B*) Line graph showing a positive correlation between wildtype transcript abundance and gene average 3′-UTR length for the X-chromosome (green) and the autosomes (grey). The genes are binned into equally sized groups based on their 3′-UTR length. (*C*) Mean values of average 3′-UTR length per chromosome arm. P-values between the X-chromosome and the 4^th^ chromosome to chromosome arm 3L (the least unlike autosome arm) are indicated. (*D, E, F*) Boxplots showing log_2_ fold change of transcript abundance in *Set2*^*1*^ versus wildtype (*D*), *ΔHisC; 12xH3K36R* versus *ΔHisC; 12x*^*H3K36K*^ (wt) (*E*), and *ΔH3*.*3B; ΔH3*.*3A* versus wildtype (*F*) with genes binned into equally sized groups based on their 3′UTR lengths. Error bars in (*A-C*) represent 95% confidence intervals. Significant differences in (*C - F*) were determined by Unpaired Two-Sample Wilcoxon Tests.

### Ash1 and NSD together maintain balanced transcriptional output from the 4^th^ chromosome

Immunostaining of polytene chromosomes and ChIP indicate that the fourth chromosome is enriched in both H3K36me2 (Supplementary Fig. S1A, C) and H3K36me3 (Fig. 1, Supplementary Fig. S1C). Furthermore, the 4^th^ chromosome genes have longer 3′-UTRs, a feature that correlated with higher immunoprecipitation with anti-H3K36me3 antibodies (Fig. 5A, C). Despite this, Set2 knock-out caused no imbalance in transcription of 4^th^ chromosome genes compared to those on autosomes.

In contrast, the *ash1*^*22*^*/ash1*^*9011*^ and the *NSD*^*ds46*^ mutants displayed a small but significant imbalance in the transcriptional output from the 4^th^ chromosome (Figs 6A, B). The effect was markedly more pronounced in flies that lacked both *ash1* and *NSD* functions (*ash1*^*22*^ *NSD*^*ds46*^*/ash1*^*9011*^ *NSD* ^*ds46*^) (Fig. 6C). By dividing the genes into housekeeping and non-housekeeping classes we found that the bulk of the change corresponded to imbalanced transcriptional output of the non-housekeeping genes (Fig. 6D). Interestingly, the magnitude of the drop in the relative transcript abundance of the non-housekeeping genes and the stronger effect on this class of genes resembled those observed in *Pof* mutants, the key component of the regulatory mechanism specific to chromosome 4 (Stenberg et al. 2009; Lundberg et al. 2013b). We conclude that Ash1 and NSD together are required for balanced transcriptional output from chromosome 4. To test if the observed reduction in *ash1*^*22*^ *NSD*^*ds46*^*/ash1*^*9011*^ *NSD* ^*ds46*^ is a consequence of impaired methylation of H3K36, we included *ΔHisC; 12x*^*H3K36R*^ for comparison. Importantly, the expression levels from chromosome 4 dropped in the H3K36R histone replacement mutants, *ΔHisC; 12x*^*H3K36R*^, by a similar extent as that in the *ash1 NSD* double mutant (Fig. 6E). Taken together these observations argue that methylation of H3K36 by both Ash1 and NSD is required to balance transcriptional output of the 4^th^ chromosome.

**Figure 6.**
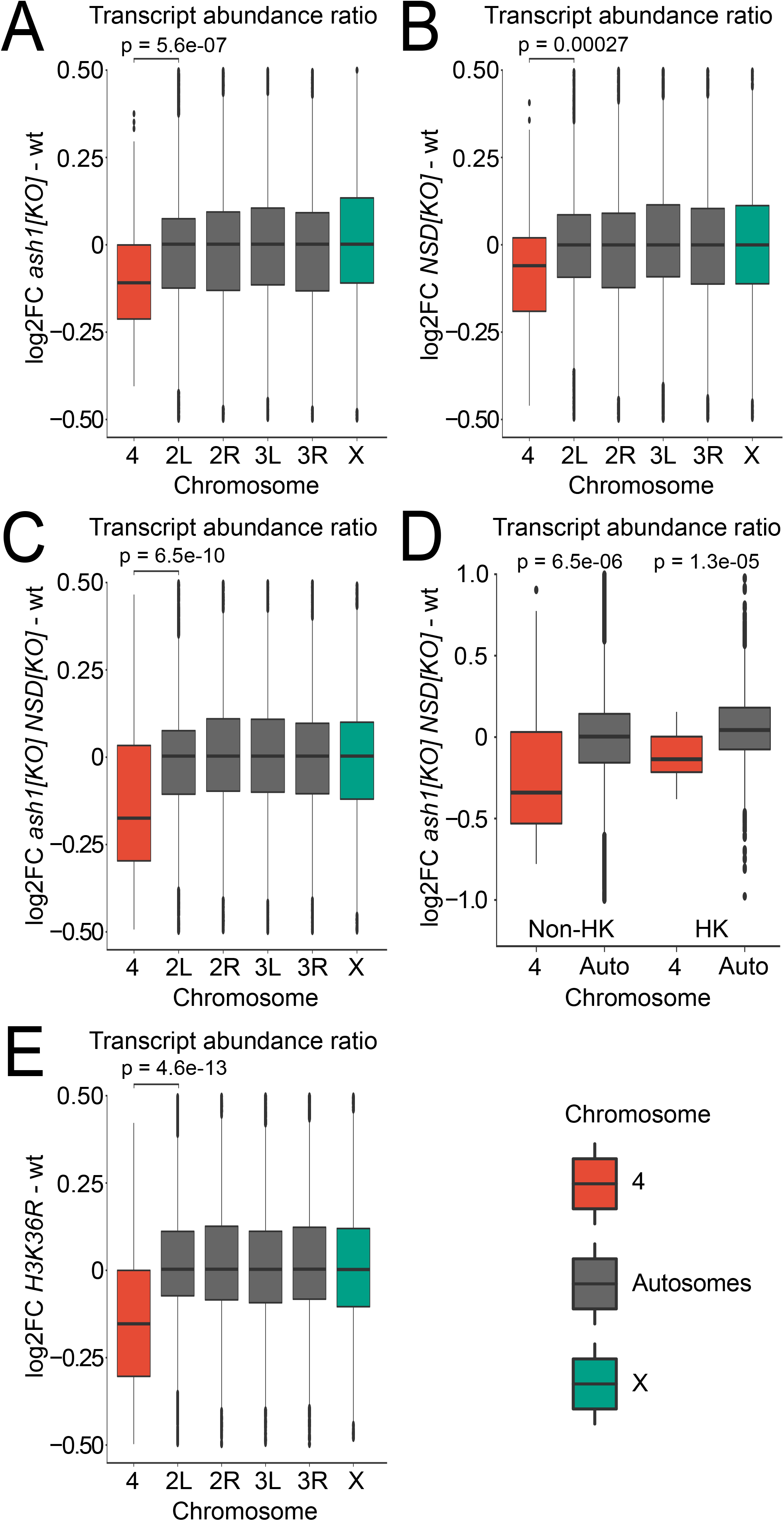
*Ash1* and *NSD* show additive effects and are required for balanced transcription output of genes on chromosome 4. Boxplot showing chromosome-specific transcript abundance ratios (log_2_ fold change) compared to wildtype for chromosome 4 (red), autosomes (grey) and the X-chromosome (green). (*A*) In *ash1*^*22*^*/ash1*^*9011*^ the transcript abundance from chromosome 4 is reduced by 8.0% compared to the main autosome arms. (*B*) In *NSD*^*ds46*^ the transcript abundance from chromosome 4 is reduced by 5.1% compared to the main autosome arms. (*C*) In *ash1*^*22*^ *NSD*^*ds46*^*/ash1*^*9011*^ *NSD* ^*ds46*^, transcript abundance is reduced by 14.9% compared to the main autosome arms. (*D*) Boxplot showing chromosome 4 transcript abundance change in *ash1*^*22*^ *NSD*^*ds46*^*/ash1*^*9011*^ *NSD* ^*ds46*^ compared to wildtype. The genes are grouped as non-housekeeping (left) and housekeeping genes (right). In *ash1*^*22*^ *NSD*^*ds46*^*/ash1*^*9011*^ *NSD* ^*ds46*^, the expression output is reduced by 21.2% in non-housekeeping genes, which is significantly greater than the 11.7% reduction observed in housekeeping genes. (*E*) Chromosome-wide transcript abundance ratios (log_2_ fold change) in *ΔHisC; 12x*^*H3K36R*^ compared to *ΔHisC; 12x*^*H3K36K*^ (wt). H3K36R causes a reduction on chromosome 4 transcript abundance compared to the main chromosome arm by 10.7%. The statistical significance was determined by an Unpaired Two-Sample Wilcoxon Test. In (*A, B, C, E*), chromosome 4 is compared to chromosome arm 2L and in (*D*), chromosome 4 is compared to all autosome genes located on chromosomes 2 and 3.

## Discussion

Two main conclusions can be drawn from the observations presented here. First, transcriptome profiling of Set2 deficient animals indicates that this methyltransferase is required for the balanced transcriptional output from the single male X-chromosome. However, the comparison with mutants where the major histone H3 isoform is replaced with a variant that carries arginine instead of lysine 36, provides evidence that the process does not involve methylation of H3K36. Second, the balanced transcriptional output from the 4^th^ chromosome requires intact H3K36 and depends on the additive functions of NSD and the Trithorax group protein Ash1.

The latter conclusion was unexpected. We and others have previously shown that Set2 is responsible for bulk of H3K36me3 (Larschan et al. 2007; Dorafshan et al. 2019). Consistently, we see an almost complete loss of H3K36me3 signal on polytene chromosomes in *Set2*^*1*^ mutants. In contrast, the loss of Set2 causes no major changes in the bulk H3K36me2 and H3K36me1 (Dorafshan et al. 2019). We therefore speculate that Ash1 and NSD stimulate transcription of the 4^th^ chromosome genes by di-methylation of H3K36. On polytene chromosomes, H3K36me2 appears enriched in pericentric heterochromatin. These regions (enriched in H3K36me2) coincide with the regions strongly enriched in H3K9me3 and HP1a (James et al. 1989; Figueiredo et al. 2012). We have previously shown that HP1a and POF together are involved in the chromosome-specific regulation of the 4^th^ chromosome (Johansson et al. 2007a; Johansson et al. 2007b; Lundberg et al. 2013b). The reduced transcriptional output from the 4^th^ chromosome upon the loss of Ash1 and NSD suggests that di-methylated H3K36 may counteract methylation of H3K9 and/or HP1a binding and repression, or that di-methylated H3K36 somehow facilitates POF functions.

The 4^th^ chromosome is enriched in repetitive DNA, and is therefore the *Drosophila* chromosome most similar in genetic composition to the human genome. In contrast to *Drosophila*, human NSD proteins are essential for proper development, and haploinsufficiencies in *NSD1* and *NSD2* have been linked to Sotos and Wolf-Hirschhorn genetic syndromes, respectively (Kurotaki et al. 2002). It is tempting to speculate that the stimulatory effect of NSD proteins may be mechanistically similar but quantitatively more important for mammalian genes embedded in the repeat-rich genome. The *Drosophila* 4^th^ chromosome displays a high and unusual tolerance to dosage differences and mis- expression (Hochman 1976; Johansson et al. 2007a; Johansson et al. 2007b; Stenberg et al. 2009; Johansson et al. 2012). Therefore, the decreased transcriptional output from the 4^th^ chromosome of the *NSD* mutants is not expected to have severe physiological consequences.

Paradoxically, our observations indicate that H3K36me3 is less abundant on the male X-chromosome while it is enriched on chromosome 4, yet it is not directly involved in either chromosome-specific regulatory system, suggesting that the differences represent a consequence rather than a cause of these chromosomes-specific functions. How could we explain the observed differences in H3K36me3 levels? Although still a matter of discussion and debate (Kuroda et al. 2016), it has been proposed that transcription elongation efficiency is higher on the male X-chromosome compared to autosomes (Larschan et al. 2011; Prabhakaran and Kelley 2012). Likewise, we have previously shown that there is a significant reduction of the transcription elongation efficiency (elongation density index) on the 4^th^ chromosome compared to the other autosomes (Johansson et al. 2012). Considering that Set2 travels with RNA polymerase II, we speculate that the faster elongation speeds on the dosage-compensated X-chromosome, leave less time for Set2 to add three methyl groups to H3K36 resulting in the overall lower H3K36me3 levels. Conversely, the slower elongation on the 4^th^ chromosome leads to more extensive H3K36 tri-methylation. H3K36me3 is associated to, and correlates with, gene expression; i.e., higher levels of H3K36me3 within genes correlate with higher transcription output. We have also shown that higher levels of H3K36me3 correlate with gene length and the length of the 3′-UTRs. We were intrigued to see that both chromosomes with specific regulatory systems have a significantly increased average length of their genes’ 3′-UTRs. Both for the X-chromosome and the autosomes, the H3K36me3 levels positively correlate with the length of 3′-UTRs, but for the X-chromosome, the overall levels of H3K36me3 are lower. H3K36me3 levels are higher on the 4^th^ chromosome compared to the other autosomes which is in line with the increased length of 3′-UTRs on this chromosome. In fact, despite the 4^th^ chromosome’s heterochromatic nature, the average level of gene expression of chromosome 4 is comparable to, or even higher than, that of genes on other chromosomes (Haddrill et al. 2008; Johansson et al. 2012).

Finally, our observations indicate that the loss of *Set2* impairs the MSL complex mediated dosage compensation while the H3.2K36R replacement does not. We hypothesise that Set2 exerts its functions by methylation of (as yet unknown) non-histone targets similar to what has recently been suggested for Ash1 (Dorafshan et al. 2019). How this contributes to dosage compensation at the mechanistic level remains to be discovered. Nevertheless, our findings imply that models that involve the binding of MSL complex to tri-methylated H3K36 need to be revised.

## Materials and methods

### Fly strains and genetic crosses

All fly strains were cultivated and crossed at 25°C in vials containing potato mash-yeast-agar. For the RNA preparations, brains from male third instar larvae were collected. Oregon R flies were used as wildtype in all experiments except in the histone preplacement comparison as described below. To generate the *ash1* mutant larvae, *ash1*^*22*^ flies (*w/+; ash1*^*22*^, *P[w*^*+mW*.*hs*^*=FRT(w*^*hs*^*) 2A]/TM3, Ser e P[w*^*+mC*^ *ActGFP]*) (Dorafshan et al. 2019) were crossed with *w*^*1118*^; *Df(3L)Exel9011/TM3, Ser e P[w*^*+mC*^ *ActGFP]*, hereafter *ash1*^*9011*^ (Dorafshan et al. 2019) and the *ash1*^*22*^*/ash1*^*9011*^ trans-heterozygote first instar larvae were isolated, based on the lack of a GFP signal under fluorescent stereomicroscopy, and grown to the third instar larvae at 25°C. The CRISPR/Cas9 generated *NSD* loss of function allele (*NSD*^*ds46*^) has previously been described (Dorafshan et al. 2019) and the *ash1 NSD* double mutant larvae (*ash1*^*22*^ *NSD*^*ds46*^*/ash1*^*9011*^ *NSD* ^*ds46*^), were generated as described previously (Dorafshan et al. 2019). To obtain *Set2* mutant larvae, we used the *Set2*^*1*^ loss of function allele (Larschan et al. 2011). Progeny from *y*^*1*^ *w*^*67c23*^ *Set2*^*1*^/*FM7c, P[GAL4-Kr*.*C]DC1 P[UAS-GFP*.*S65T]DC5* were screened under a fluorescent stereomicroscope and non-GFP larvae (homozygous and hemizygous for *Set2*^*1*^) were transferred to separate vials. To generate *ΔHisC; 12x*^*H3K36R*^ and *ΔHisC; 12x*^*H3K36K*^ larvae, the fly strain *w; UAS-2*x*YFP ΔHisC/CyO, P[ftz-lacZ]* was crossed with either *y w; elav-Gal4, ΔHisC/CyO; VK33[H3K36Rx12]/TM6B,Tb* (to supplement the progeny with the transgenic mutant Histone cluster where Lys36 residue of *His3*.*2* was replaced by Arg), or *y w; elav-Gal4, ΔHisC/CyO; VK33[H3K36Kx12]/TM6B,Tb* (to supplement the progeny with the transgenic wildtype Histone cluster) (McKay et al. 2015). The larvae with the correct genotype were then selected based on the presence of a GFP signal (hence homozygous for deletion of endogenous Histone cluster (*ΔHisC*)), and an absence of the *Tb* marker (hence presence of the transgenic Histone cluster). The *ΔH3*.*3B; ΔH3*.*3A* mutant larvae (Hodl and Basler 2009) were isolated based on the lack of a GFP signal under fluorescent stereomicroscopy in the progeny from the cross *w, ΔHis3*.*3B, hsp-Flp; Df(2L)His3*.*3A/CyO, P[w*^*+mC*^ *ActGFP]JMR1 × w, ΔHis3*.*3B, hsp-Flp/Y; Df(2L)His3*.*3A/CyO, P[w*^*+mC*^ *ActGFP]JMR1*.

### Immunostaining of polytene chromosomes

Immunostaining of polytene chromosomes was done essentially as described previously (Johansson et al. 2012; Lundberg et al. 2013a). The antibodies used in the protocol are listed in Supplementary Table S1. Preparations were analysed using a Zeiss Axiophot microscope (Plan Apochromat 40x/0.95 objective) equipped with a KAPPA DX20C CCD camera and with Zeiss Apotome Microscope (Plan Apochromat 63x/1.40 oil DIC M27 objective, filters set: 63HE for red channel, 38HE for green channel and 49 for DAPI) equipped with AxioCam MR R3 camera. For comparisons of targeting between different genotypes, the protocol was run in parallel and nuclei with clear cytology were chosen on the basis of DAPI staining and photographed. At least 20 nuclei per slide were used in these comparisons and at least five slides per genotype. Images were processed with ZenPro software (v2.3, Zeiss).

### Western blot

Total tissue extracts were prepared from hand-dissected brains and imaginal discs of third instar larvae. For analysis of the H3K36 methylation, protein extracts were loaded on a 15% SDS–PAGE gel and blotted to a PVDF membrane for 60 minutes at 200 mA. Primary and secondary antibodies were diluted in 1xPBS, 1% BSA, 0.05% Tween-20. The antibodies used in the protocol are listed in Supplementary Table S1.

### Library Preparation

For each sample, five male third instar larvae brains were homogenised and RNA was purified using the Directzol microprep kit (Zymo research) and using the standard protocol with on-column DNase treatment. RNA quality was determined with a Fragment Analyser 5200 (Agilent Technologies, Inc.) using the DNF-471 Standard Sensitivity RNA reagent kit. Total-RNA libraries were synthesised using the Ovation RNA-Seq system for *Drosophila* (NuGEN). Standard protocol procedure with integrated DNase treatment was used to make the sequencing libraries. Fragmentation was performed using a Covaris E220 Focused-ultrasonicator with the recommended settings for 200 base pair target length. The library quality was evaluated on a Fragment Analyser using the DNF-920 DNA reagent kit.

### Sequencing and data analyses

In the first round, a total of five samples per genotype were sequenced (Oregon R, *Set2*^*1*^, *ash1*^*22*^*/ash1*^*9011*^, *NSD*^*ds46*^, *ash1*^*22*^ *NSD*^*ds46*^*/ash1*^*9011*^ *NSD* ^*ds46*^, *ΔHisC; 12x*^*H3K36R*^ and *ΔHisC; 12x*^*H3K36K*^) on Illumina HiSeq2500 (Illumina Cambridge Ltd.) by Science for Life Laboratory, Stockholm and 125 bp paired-end reads were obtained. In the second round, seven samples of *ΔH3*.*3B; ΔH3*.*3A* and an additional six Oregon R samples were sequenced at Science for Life Laboratory, Stockholm using Illumina NovaSeq SP (Illumina Cambridge Ltd.) with a paired-end read length of 150 bp. Reads from both rounds were mapped to the *D. melanogaster* genome version BDGP6 using STAR v2.5.3 (Dobin et al. 2013) with default settings. Read counts were obtained with featureCounts v1.5.1 with default settings (Liao et al. 2014). Gene abundances were calculated using StringTie v1.3.3 (Pertea et al. 2015). Replicate 3 of *Set2*^*1*^ and replicate 5 of *ΔHisC; 12x*^*H3K36K*^ did not pass the quality control in RSeQC (Wang et al. 2012) and were therefore excluded in the downstream analysis. DESeq2 v1.18 (Love MI *et al*. 2014) was used for differential expression analysis using default parameters and normal log fold change shrinkage.

### Distance to High Affinity Sites (HAS) and MSL gene enrichments

Genomic locations for 188 previously published High Affinity Sites, compiled by (Philip and Stenberg 2013), originating from (Alekseyenko et al. 2008) (Straub et al. 2008) were used in the analysis. The distance to the closest HAS was calculated for each gene on the X-chromosome. The gene enrichment data for the MSL complex component MSL1 were from (Philip and Stenberg 2013). The binding data were subdivided into five equally sized bins based on the average MSL1 enrichment, bin 1 comprising the genes with the lowest MSL1 enrichments, up to bin 5 which comprised genes with the highest MSL1 enrichments.

### Classification of genes as early or late replicating

Genes were classified as early or late replicating as previously described (Kim et al. 2018b). The data on early and late replicating genomic domains were kindly provided by David MacAlpine (Lubelsky et al. 2014) and the coordinates were converted from *Drosophila* genome release 5 to release 6 using flybase.org’s online coordinate converter. The annotating tool from BEDTools (Quinlan and Hall 2010) was used to calculate the overlap between genes and replication domains. Genes were classified as early or late replicating if the entire transcript was within a domain classified as early or late replicating in the male S2 cell line.

### Classification of genes as housekeeping or non-housekeeping

Genes with >6 as the expression level in all 12 FlyAtlas-specified tissue types (Chintapalli et al. 2007) are defined as housekeeping genes and genes with >6 expression levels in 11 or fewer tissue types are here defined as non-housekeeping genes or differentially expressed genes.

### Wildtype gene expression

The average Transcript per million (TPM) score for wildtype samples (Oregon R) was calculated for all genes and subsequently binned according to Flybase RNA-Seq convention (Thurmond et al. 2019). The bins are 0-0 TPM, 1-3 TPM, 4-10 TPM, 11-25 TPM, 26-50 TPM, 51-100 TPM, 101-1000 TPM and >1000 TPM for Unexpressed, Very low expression, Low expression, Moderate expression, Moderately high expression, High expression, Very high expression and Extremely high expression, respectively.

### Datasets and calculated gene enrichments

Raw and processed sequencing data generated were deposited to NCBI GEO under accession GSE166934. ChIP-chip data sets were obtained from GSE23457, GSE23458 and GSE32793. ChIP-seq data sets were obtained from GSE127177 (Prayitno et al. 2019) and, in processed form, kindly provided by Tamás Schauer and Peter Becker. RNA-seq data from the temperature sensitive *E(z)*^*61*^ cell line were obtained from GSE61307 (Lee et al. 2015) The *E(z)*^*61*^ cell line was classified as male based on the expression of *roX1, roX2, msl2* and *traF*. Enrichment scores for the ChIP-chip experiments is a log_2_ ratio of ChIP over input. A custom script was used to calculate the top 50 percent of the total exonic region for each gene. For ChIP-seq the Homer suite v4.11 (Heinz et al. 2010) was used to obtain H3K36me3 peaks over input. The peak score used for downstream analysis is defined as position adjusted reads from initial peak region. Only peaks originating in exons were used when calculating the chromosomal average.

### Bioinformatics and plotting of the figures

All calculations after the compilation of the raw counts table were performed using R (R_Core_Team 2016) and plots were generated using the ggplot2 R package (Wickham 2009).

## Acknowledgments

We thank Tamás Schauer and Peter Becker for sharing H3K36me3 ChIP-seq data; Dirk Schübeler for the NSD antibody; Mitzi Kuroda, Gregory Matera and Konrad Basler for fly strains. We also thank the Science for Life Laboratory, Stockholm, Sweden; the National Genomics Infrastructure (NGI), Stockholm, Sweden; and UPPMAX, Uppsala, Sweden, for assistance with the RNA sequencing and for providing the computational infrastructure. This work was supported by grants from the Swedish Research Council (2016-03306 to J.L. and 2017-03918 to Y.B.S.) Knut and Alice Wallenberg Stiftelse (2014.0018 to J.L. and Y.B.S.), Nils Erik Holmstens Forskningsstiftelse (to Y.B.S.) and Swedish Cancer Foundation (CAN 2017/342 to J.L.).

## Author contributions

JL and HL conceived the project. HL performed the RNA-seq experiments and all bioinformatics. AG and JL performed the immunostainings and AG performed the Western blots. HL and ED performed the fly genetics. HL, AG, ED, YBS and JL together analysed the data. HL, YBS and JL wrote the manuscript with contributions from all authors.

## Figure legends Supplementary figures

**Supplementary Figure S1.**
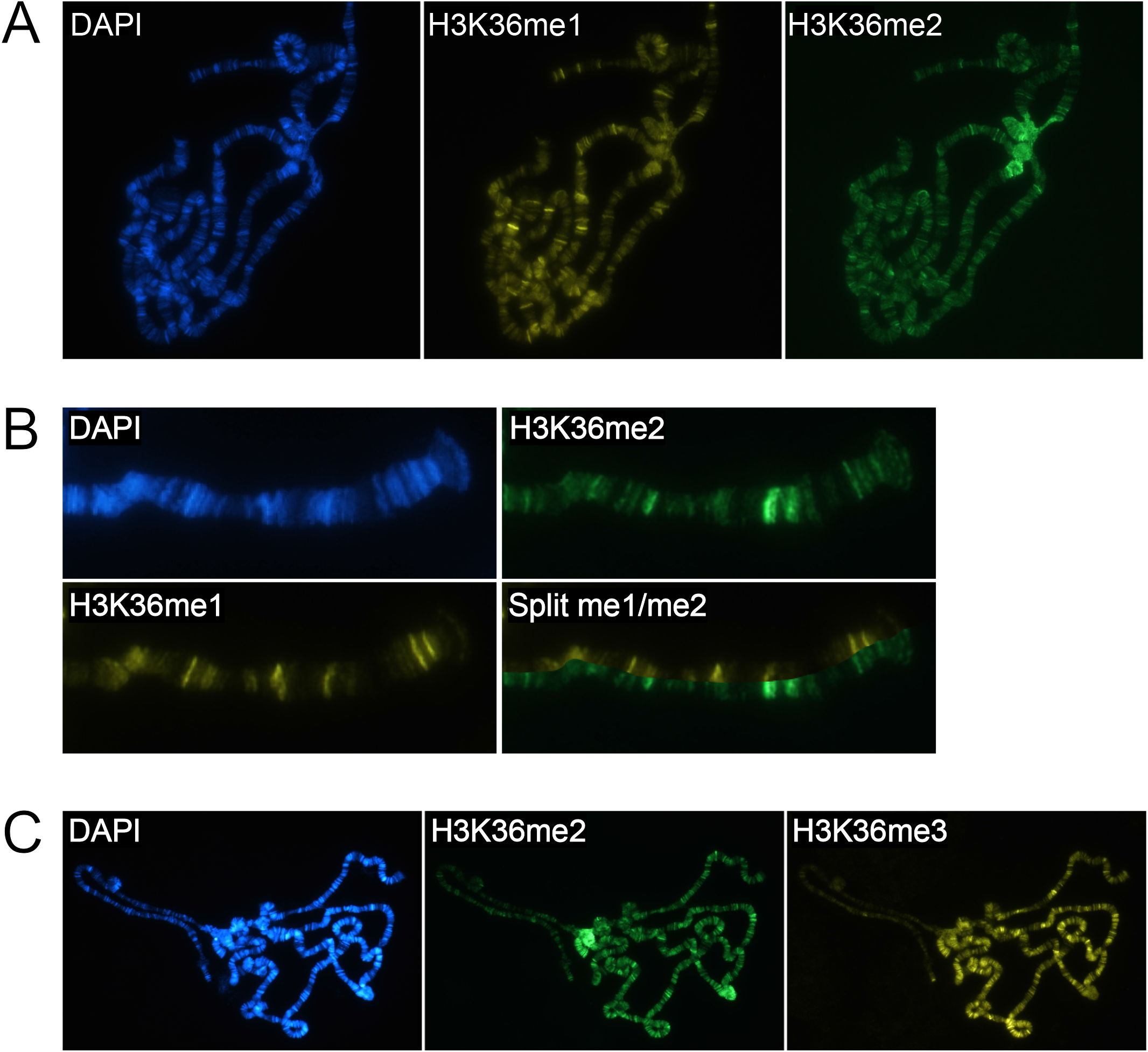
Differential enrichments of methylated H3K36. (*A*) Immunostaining of third instar larvae polytene chromosome showing H3K36me1 (yellow), H3K36me2 (green) and DAPI staining of DNA in blue. Note the accumulation of H3K36me2 in pericentromeric heterochromatin. (*B*) The H3K36me2 (green) staining shows only minor overlaps with H3K36me1 (yellow). (*C*) Immunostaining of a male third instar larvae polytene chromosome showing H3K36me2 (green), H3K36me3 (yellow) and DAPI staining of DNA in blue. Note the accumulation of H3K36me2 in pericentromeric heterochromatin, H3K36me3 on the 4^th^ chromosome and the reduced amount of H3K36me3 on the X-chromosome.

**Supplementary Figure S2.**
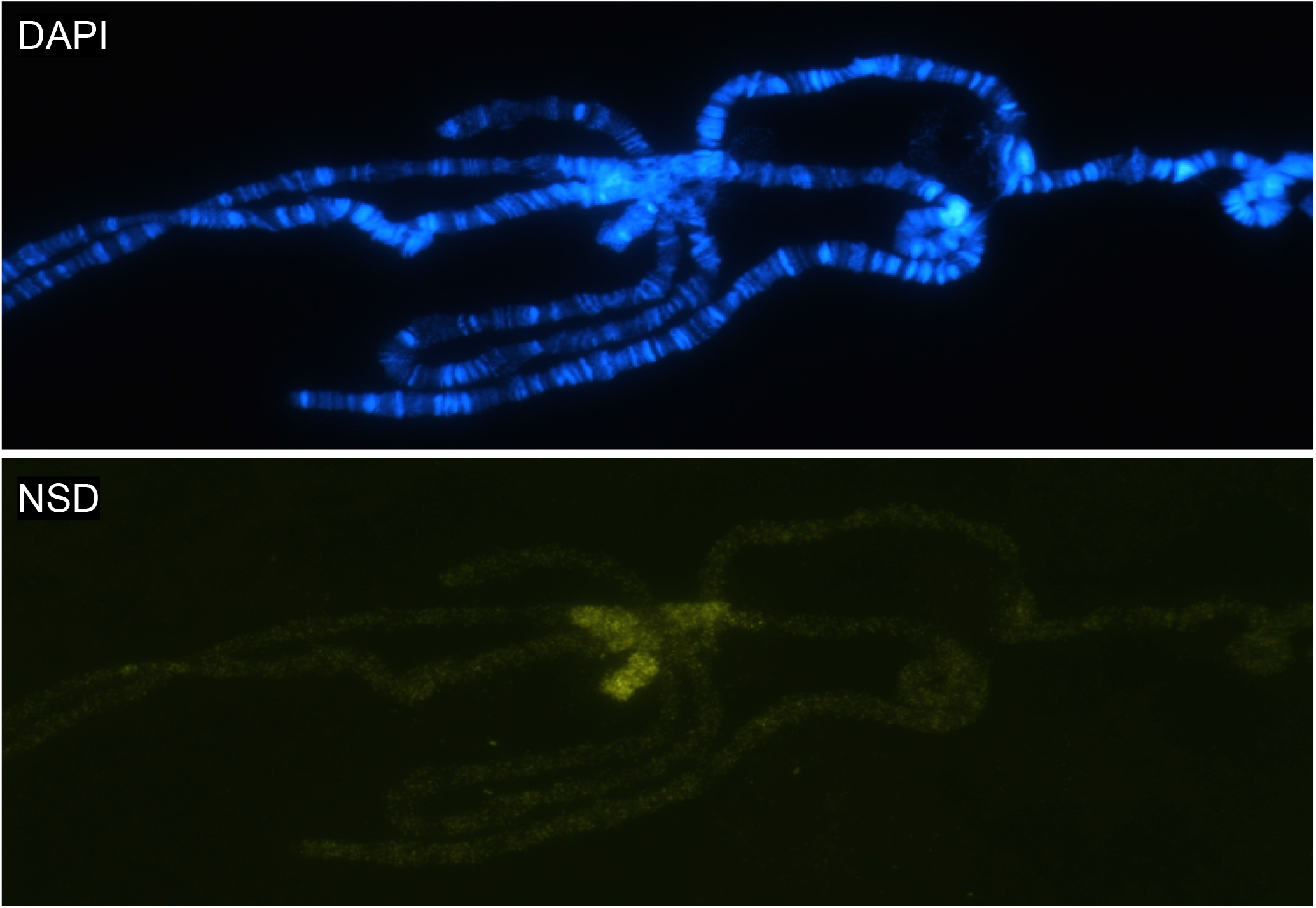
NSD targets pericentric heterochromatin. Immunostaining of third instar larvae polytene chromosome showing NSD (yellow), and DAPI staining of DNA in blue. Note the accumulation of NSD in pericentromeric heterochromatin and region 2L:31, similar to typical enrichments of HP1a.

**Supplementary Figure S3.**
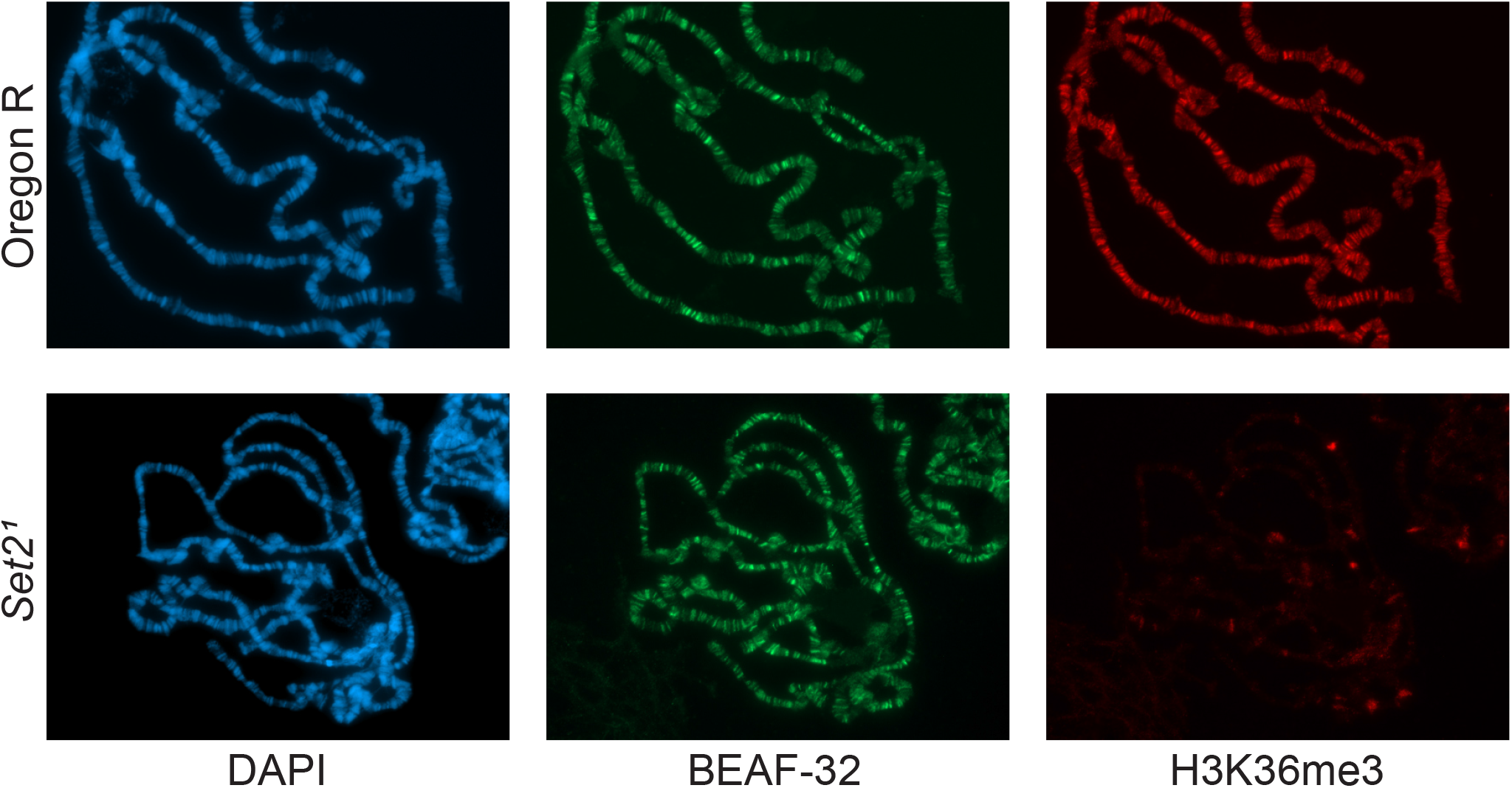
H3K36me3 is lost in *Set2*^*1*^ mutants. Staining of H3K36me3 in salivary glands for wildtype (top row) and *Set2*^*1*^ (bottom row). Staining of BEAF-32 serves as an internal control.

**Supplementary Figure S4.**
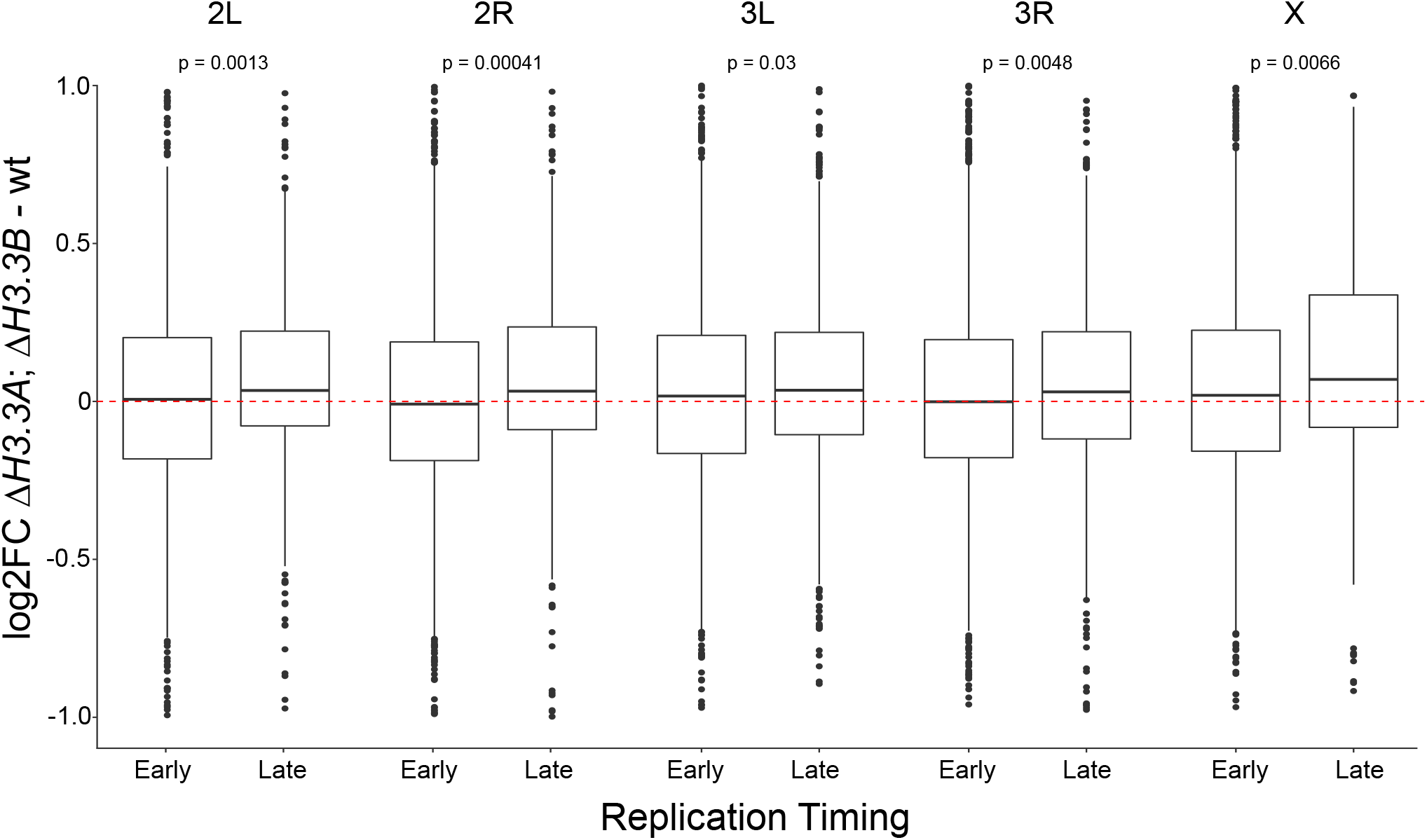
Boxplot showing log_2_ fold change of transcript abundance in *ΔH3*.*3B; ΔH3*.*3A* compared to wildtype with genes divided by chromosome arms, and into early or late replicating. Note the small but significant relative increase of expression output from late replicating genes. The statistical significance was determined by an Unpaired Two-Sample Wilcoxon Test.

**Supplementary Table S1:**
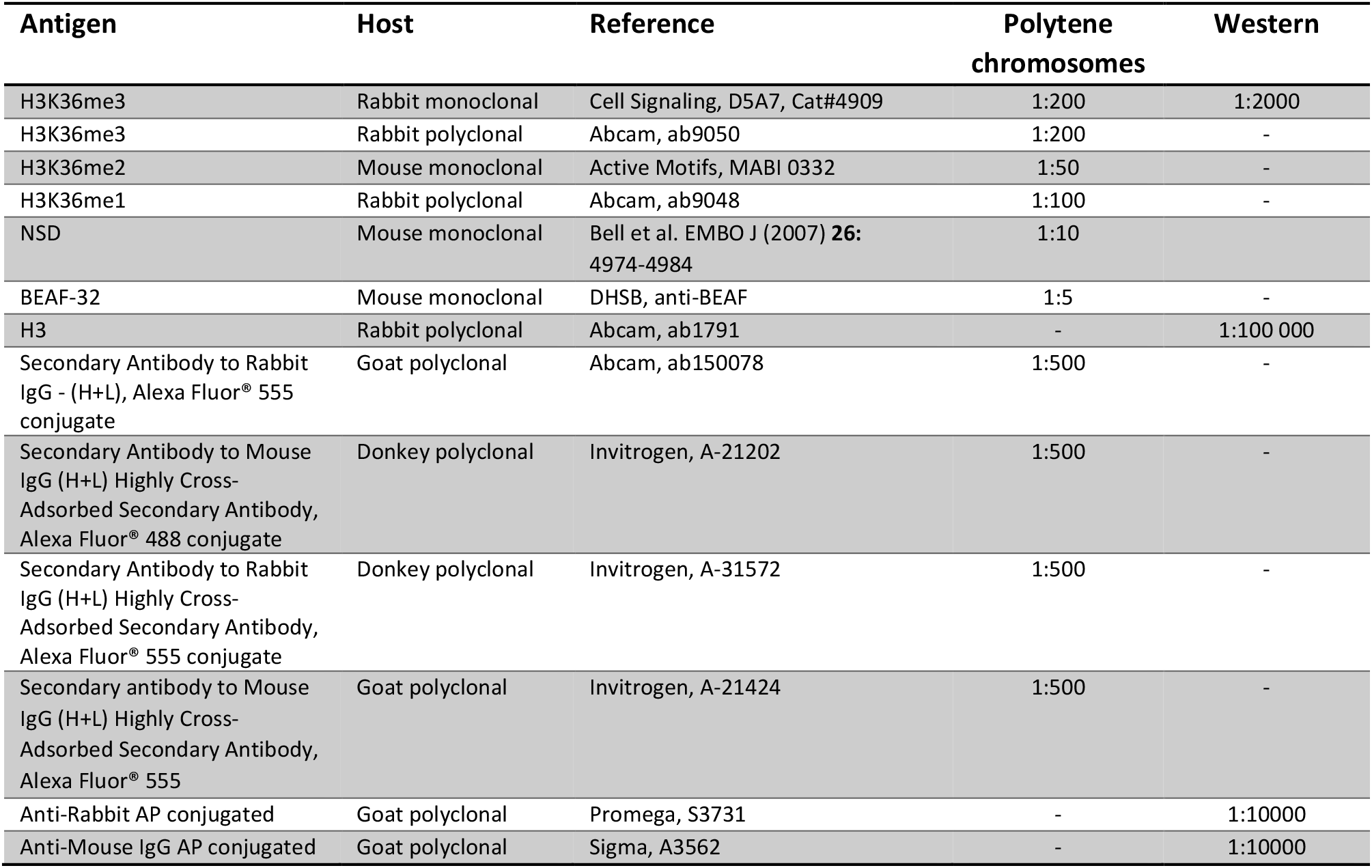
List of antibodies.

